# Rational design of peak calling parameters for TIP-seq based on pA-Tn5 insertion patterns improves predictive power

**DOI:** 10.1101/2024.10.08.617149

**Authors:** Thomas Roberts, Hiranyamaya Dash, Teemu K. E. Rönkkö, Maria Weinert, Nathan Skene

## Abstract

Epigenomic profiling provides insights into the regulatory mechanisms that govern gene expression. At a fundamental level, these mechanisms are determined by proteins that bind the DNA or modify the chromatin. Techniques such as ChIP-seq and CUT&Tag have been instrumental in mapping the binding sites of such proteins across the genome. Recent advances have led to the development of TIP-seq, a highly sensitive method devised to increase the number of unique reads per sample. Its design results in novel library features, which have not yet been explored with comparative analytics. Through the extensive assessment of bioinformatics tools and parameters we have developed an analysis pipeline that is ideally suited for TIP-seq data, including linear deduplication, read prioritisation and read shifting. Using transcription factor binding profiles (TFs), we show that our optimised pipeline greatly reduces the width of peaks to below 50% and more precisely aligns the peak summit with known motifs. A tutorial of the optimised peak calling is available on GitHub at https://github.com/neurogenomics/peak_calling_tutorial.git. Our methodological advancement substantially improves TIP-seq data quality, and the thoughtful design of analysis parameters is widely applicable to all pA-Tn5 based profiling assays.

## Introduction

New methods like Cleavage Under Target and Tagmentation (CUT&Tag) and related techniques leverage the action of the Tn5 transposase to profile protein binding to DNA including in single cells (Bartlett et al., 2021; Bartosovic et al., 2021; Carter et al., 2019; Janssens et al., 2024; Kaya-Okur et al., 2019; Meers et al., 2023; Wang et al., 2019; Wu et al., 2021; Xiong et al., 2024; Zhu et al., 2021). Specifically, protein A-tagged Tn5 (pA-Tn5) is directed to chromatin-bound antibodies, where it simultaneously cuts and inserts adapters at target regions. As a result, these methods require lower cell input and produce less background than Chromatin Immunoprecipitation followed by sequencing (ChIP-seq), which ENCODE used extensively to generate maps of transcription factor (TF) binding and histone marks (Landt et al., 2012). The classic CUT&Tag protocol inserts forward and reverse PCR adapters (Kaya-Okur et al., 2019), which is suboptimal as successful amplification requires both adapters to be inserted in the correct orientations. Tip-seq (Bartlett et al., 2021) circumvented that problem through insertion of a single adapter containing a T7 promoter, followed by linear amplification of antibody-bound regions through an RNA intermediate, which substantially increases the sensitivity of the assay. The second PCR adapter is introduced during an unspecific tagmentation to complete library construction. Later, Nano-CT (Bartosovic & Castelo-Branco, 2023) has adopted that strategy albeit without the RNA intermediate. In the latest versions of CUT&Tag-based protocols, namely NTT-seq and uCoTarget, the forward and reverse PCR adapters are inserted consecutively by Tn5 bound to primary and secondary antibodies, respectively (Stuart et al., 2023; Xiong et al., 2024), which likely increase the number of amplifiable fragments. With respect to data analysis, these enhanced protocols share a unique feature. The 5’ end of the first adapter is spatially the closest estimate for protein binding to the target DNA due to the sequential nature of adapter insertion.

To identify target DNA several algorithms have been developed that determine regions with an enrichment of sequencing reads against the background, a process referred to as peak calling. The Python program MACS was originally developed for ChIP-seq but has evolved into a general peak caller that can be applied to any DNA enrichment assay (Zhang et al., 2008). MACS employs a dynamic Poisson distribution to identify genomic regions with significantly higher read (tag) densities compared to the background. In this context, a Poisson distribution implies that the occurrence of reads in each region follows a Poisson process with rate parameter l, which refers to the expected read number across a given region. This parameter is calculated in a region-specific manner to account for local biases. However, if the background is dominated by zeroes, even regions with small levels of read enrichment may be called significant.

Low background signal compared to ChIP-seq is a feature of Tn5-based methods, like TIP-seq. This has motivated the development of peak callers like SEACR (Sparse Enrichment Analysis for CUT&RUN), which has been proposed to counter the oversensitivity of MACS on low-background DNA enrichment assays (Meers et al., 2019). First, SEACR identifies genomic regions (or blocks) with non-zero signal. The program then filters out all blocks that have a signal value below a global threshold, derived from either IgG controls or user-supplied (not recommended). Although SEACR may seem appropriate for methods like TIP-seq, recent work has demonstrated the superiority of MACS, at least for narrow marks like TFs and some histones (Yashar et al., 2022). An additional advantage of MACS over SEACR is the identification of peak summits, which are the exact nucleotides where signal enrichment is highest and serve as a proxy for binding sites. This has led to the adoption of MACS as industry standard for peak calling.

Given the fast pace of targeted epigenetics methods development, little work has been dedicated to optimising data analysis. Instead, already established pipelines, including MACS peak calling, are considered suitable with default Chip-seq settings (Zheng et al., 2020). As a consequence, little consideration is given to select parameters that reflect the unique properties of the data and reporting of chosen settings is often poor or misleading (Bartosovic et al., 2021; Lyu et al., 2022). The effect of research dedicated to data analysis is unknown but will likely result in further improvements in data quality, which is of interest to the whole field.

## Methods

### Retrieving publicly available TIP-seq data

We obtained TIP-seq data (raw reads) generated by Bartlett et al. (2021) from NCBI GEO under accession number GSE188512. The authors performed TIP-seq using 5000 HCT116 colorectal adenocarcinoma cells, targeting the TF CTCF.

### iPSC microglia differentiation

Microglia-like cells (MGL) were differentiated from human induced pluripotent stem cells (iPSCs) via primitive macrophage precursors (pMacPre) according to Haenseler et al. (2017) as previously described (Müller et al., 2023) with small variations: The iPSC line SFC841-03-01 (StemBANCC) was cultured in OXE8 medium (Vaughan-Jackson et al., 2021) on Geltrex (Gibco, Thermo Fisher Scientific, UK). For embryoid body production, 3×10^6^ iPSC were plated in one well of a 24-well Aggrewell 800 plate (STEMCELL Technologies, UK) in EB medium (OXE8, 20 ng/mL SCF (Miltenyi Biotec, UK), 50 ng/mL BMP4 (Gibco), 50 ng/mL VEGF (PeproTech, UK)). Embryoid bodies were fed daily for one week, then transferred to 2x T175 flasks containing FM7a factory medium (Vaughan-Jackson et al., 2021). For 4 weeks, factories were fed weekly with 10 mL FM7a, thereafter pMacPre producing flasks (>95% triple-positive for CD45, CD14 (both Immunotools, Germany) and CD11b (Biolegend, UK)) were fed weekly with 30 ml FM7a. pMacPre were harvested from the supernatant, plated at 20,000 cells/cm2 in 10-cm culture dishes and differentiated into MGL for 7 days in MIC10 medium (SILAC Advanced DMEM/F12, 2mM GlutaMax (both Gibco), 10 mM glucose, 0.5 mM L-lysine, 0.7 mM L-arginine, 0.00075% phenol red (all Sigma Aldrich, UK), 100 ng/mL IL-34 (PeproTech), 10 ng/mL GM-CSF (Gibco)). An equal volume of MIC10 was added after 3 days in culture.

### Cell harvest

MGL were incubated for 2 min at 37ºC in TrypLE and transferred to 2 ml 10% BSA. The remaining cells were gently scrape on ice in PBS and pooled. Cells were spun for 5 min at 500xg, resuspended in freeze medium (50% MIC10, 10% DMSO (Sigma), 10% KO-DMEM, 30% ES-FBS (both Gibco)), and cryopreserved at -80ºC. Cells were quickly thawed directly prior to the assay, spun as above and resuspended in wash buffer (20 mM HEPES pH 7.5, 150 mM NaCl, 0.5 mM Spermidine, 1x EDTA-free cOmplete Protease Inhibitor Cocktail).

### Tn5 and pA-Tn5 loading

All oligonucleotides were resuspending in nuclease-free water to a concentration of 100 mM. Adaptors were annealed by mixing equimolar amounts of ME-T7 and ME-rev or ME-B and ME-rev, denaturing at 95°C for 5 min and cooling to 20°C at a rate of 0.1°C/s. pA-Tn5 (C01070002, Diagenode) was loaded with ME-T7 adaptors and Tn5 (C01070010-20, Diagenode) was loaded with ME-B adaptors by incubating 4 ul of 50 uM annealed adaptor, 5 ul transposase, and 21 ul glycerol in 20 ul annealing buffer (0.1 M HEPES, pH7.2, 0.2 M NaCl, 0.2 mM EDTA, 0.2% TritonX-100, 20% Glycerol, 2 mM DTT)) for 1 h at room temperature. Loaded transposases were stored at -20°C for up to 1 month.

### Tip-seq

Tip-seq was performed according to Bartlett et al. (Bartlett et al., 2021). In brief, 50,000 cells were conjugated to equilibrated ConcavalinA beads (BP531, Bangs Laboratories) in wash buffer prior to incubation with primary antibody (CREB1,1:50, #9197, Cell Signalling) for 16h at 4°C in antibody buffer (Wash buffer, 0.01% digitonin (300410, Millipore), 2 mM EDTA, 1% BSA), secondary antibody (1:50, guinea pig anti-rabbit, ABIN101961, antibodies-online) for 1h in Dig-wash buffer (Wash buffer, 0.01% digitonin) and pA-Tn5/ME-T7 (1:25) for 1h in Dig300 buffer (Dig-wash buffer with a total of 300 mM NaCl). After washing DNA was tagmented (Dig-300 Buffer, 10 mM MgCl2) for 1h at 37°C and 500rpm. Cells were lysed in 4M Guanidine Hydrochloride with 16 mM EDTA and the DNA purified with AMPure XP beads (A63880, Beckman Coulter) without bead removal after elution. Gap-fill was performed with 5xTaq polymerse (M0285, New England Biolabs) for 3 min at 72°C prior to In-vitro transcription for 16-19 h at 37°C using the HiScribe T7 High Yield Synthesis kit (E2040S, New England Biolabs) including 0.3 μl RNase Inhibitor (M0314L, New England Biolabs)). RNA was purified with 2x volume of SPRI RNA binding buffer (20% PEG 8000, 2.5 M NaCl, 1 mM citrate pH 6.4). Reverse transcription was primed with random hexamers (N8080127, Thermo Fisher Scientific) and performed using SMART MMLV RT kit (639524, Takara) and dNTPs (R0192, Thermo Fisher Scientific) at 22°C for 10 mins, 42°C for 60 min, then 70°C for 10 min. RNA was degraded with 0.5 U RNase H (M0297L, New England Biolabs) for 20 min at 37°C. The second strand was primed with 2.5 ml 20 uM sss_bulk oligonucleotide at 65°C for 2 min and extended with 5xTaq polymerase for 8 min at 72°C. cDNA was purified with 2x volume of SPRI DNA binding buffer (20% PEG 8000, 2.5 M NaCl, 10 mM Tris-HCl pH 8.0, 1 mM EDTA) and tagmented with 2 ul 0.7 mM Tn5/ME-B and 2 ul 5x TAPS buffer (10 mM TAPS, 0.2 mM EDTA) for 10 min at 55°C, the complex dissociated in 4 M guanidine hydrochloride, and the cDNA purified with 2x volume of SPRI DNA binding buffer with removal of beads. PCR amplification was performed using primers P501 and N701 in NEBNext 2x Mastermix (M0541L, New England Biolabs) at 72°C 3 minutes, 95°C 30 seconds, 5 cycles of 98°C for 10 seconds, 63°C for 30 seconds, 72°C for 1 minute. Additional PCR cycles were determined as the cycle number corresponding to 1/4^th^ of the maximum sample fluorescence using a qPCR side reaction according to Buenrostro et al. (2015). The libraries were size-selected first with 0.5x volume AMPure XP beads followed by 0.85x volume of AMPure XP beads. The fragment size distribution was assessed by TapeStation HS D5000 (5067-5582, Agilent) and sequenced on Novagene 6000 (Illumina) at 10 million reads per sample resulting in ∼6,500,000 read pairs compared to ∼3,000,000 read pairs for the CTCF data from Bartlett et al. (2021) after quality control and removing linear duplicates.

### Oligonucleotides

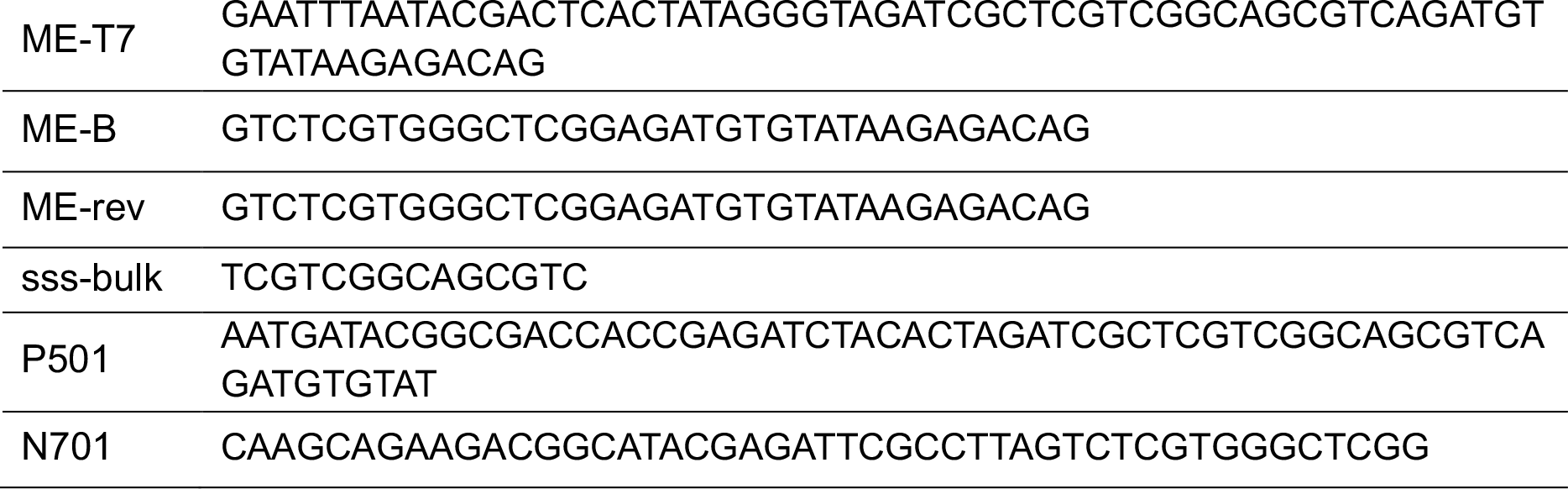

### Processing TIP-seq data with the nf-core/cutandrun pipeline and peak calling

All data was processed using the *nf-core* pipeline CUT&Run v3.2.0 (Cheshire et al., 2023). Duplicates resulting from linear amplification were removed by setting *remove_linear_duplicates* to ‘true’, a function that we recently added to the pipeline. This setting is predicated on the principle that reads with identical 5’ start sites are likely to originate from the same linear amplification event: this is a divergence from the published TIP-seq processing pipeline, which kept some of the linear duplicates. Peaks were called with MACSr, a MACS3 wrapper, with the following settings: *nomodel = TRUE, extsize = 150, shift = -75, keep_duplicates = all*, and *q-value = 0*.*01 (Hu, 2024)* using two types of BAM files: the standard paired-end BAM file generated by the CUT&Run pipeline and a filtered BAM file containing only the first mate (first read) as generated by *samtools view -hb -f 64 paired_end*.*bam > first_mate*.*bam* (H. Li et al., 2009). No controls were used for peak calling because our IgG controls had considerably fewer reads and MACS scales the larger sample to the same read count as the smaller sample, removing much of the signal.

### Motif enrichment and *De novo* motif discovery

Motifs for CTCF (MA1930.2) and CREB1 (MA0018.5) were retrieved from JASPAR (Rauluseviciute et al., 2024). Motif enrichment in peaks was calculated using the function *runAme* from the R package memes relative to background sequences (Nystrom & McKay, 2021). *runFimo* was used to find individual occurrences of motifs in peaks. We attempted to threshold using a q-value of 0.05, but the results were extremely conservative for the short CREB1 motif. This is a known issue when scanning for short motifs, where even a perfect match may not be strongly statistically significant (Grant et al., 2011). Instead, we calculated a p-value threshold adjusted for the false positive (FP) rate and average peak width as follows:

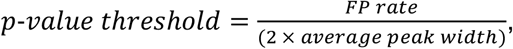

where *FP rate* was set to 0.05. This threshold adjustment is required as random motif occurrences increase with a wider search space (peak). The formula is an empirically derived heuristic that works well in practice (Grant et al., 2011). For both *runAme* and *runFimo*, a 0-order background model was generated using the letter frequencies in the primary sequences. The background model is used to produce a set of control sequences with equal length and number to the primary sequences.

The distance between each MACS3 peak summit (highest fragment pileup) and its nearest motif was calculated using the *summit_to_motif* function in the R package MotifStats, which uses *runFimo* to retrieve the motif locations. Note that the summit-to-motif distance is calculated from the centre of the motif.

STREME from the MEME suite was used to perform *De novo* motif discovery. A 2^nd^-order background model was used to generate background sequences with the same trinucleotide frequencies as the primary sequences (e.g. if the frequency of TTC in the primary sequences is 2%, this will be reflected in the background sequences). As input we used peaks that were centred on their summits and extended to fixed widths of 50 bp, 100 bp, 150 bp, 200 bp, 250 bp, and 300 bp.

## Results

When designing the analysis strategy for a new whole-genome profiling technique, the specific characteristics of the experimental workflow will influence the analysis parameters and interpretation of the sequencing reads. For example, Tip-seq was the first workflow using linear amplification of targeted DNA, which can produce linear duplicates in addition to the PCR duplicates that occur during library amplification. In paired-end sequencing, linear duplicates share the start site of the first read characterising the original tagmentation site but differ in the location of the second read, due to unspecific secondary tagging. If these duplicates are not correctly removed, an overestimation of reads will skew the data. To facilitate removal of linear duplicates, we have implemented the function *remove_linear_duplicates* (see Methods), which will aid correct pre-processing of Tip-seq and other workflows with linear amplification, like Nano-CT, as part of the NFCore CUT&Run pipeline.

We further analysed the Tip-seq workflow to develop an optimised analysis pipeline, which should improve the data quality and reliability. After binding to the target, pA-Tn5 cleaves the DNA at one of four potential sites relative to the protein of interest (Fig. 1), followed by DNA amplification and secondary tagmentation to introduce the second sequencing adaptor. We identified two key characteristics that would affect the interpretation of sequencing reads. First, amplification of the target DNA occurs downstream towards the 3’ end resulting in a 3’ amplification bias of the sequencing reads with respect to the original cut site, while the 5’ end of each read reflects the closest known location to the target (Fig 1). Second, and specific to Tip-seq and other methods that employ linear amplification, the original cut site is reflected only in the first read when performing paired-end sequencing, while the second read is produced by an unspecific tagmentation at an arbitrary location not governed by the protein of interest. We therefore reasoned that 1) centring all first reads on their 5’ end will overcome the 3’ amplification bias and 2) filtering the paired-end sequencing data to retain only first reads will improve the predictive power of Tip-seq peaks.

**Figure 1.**
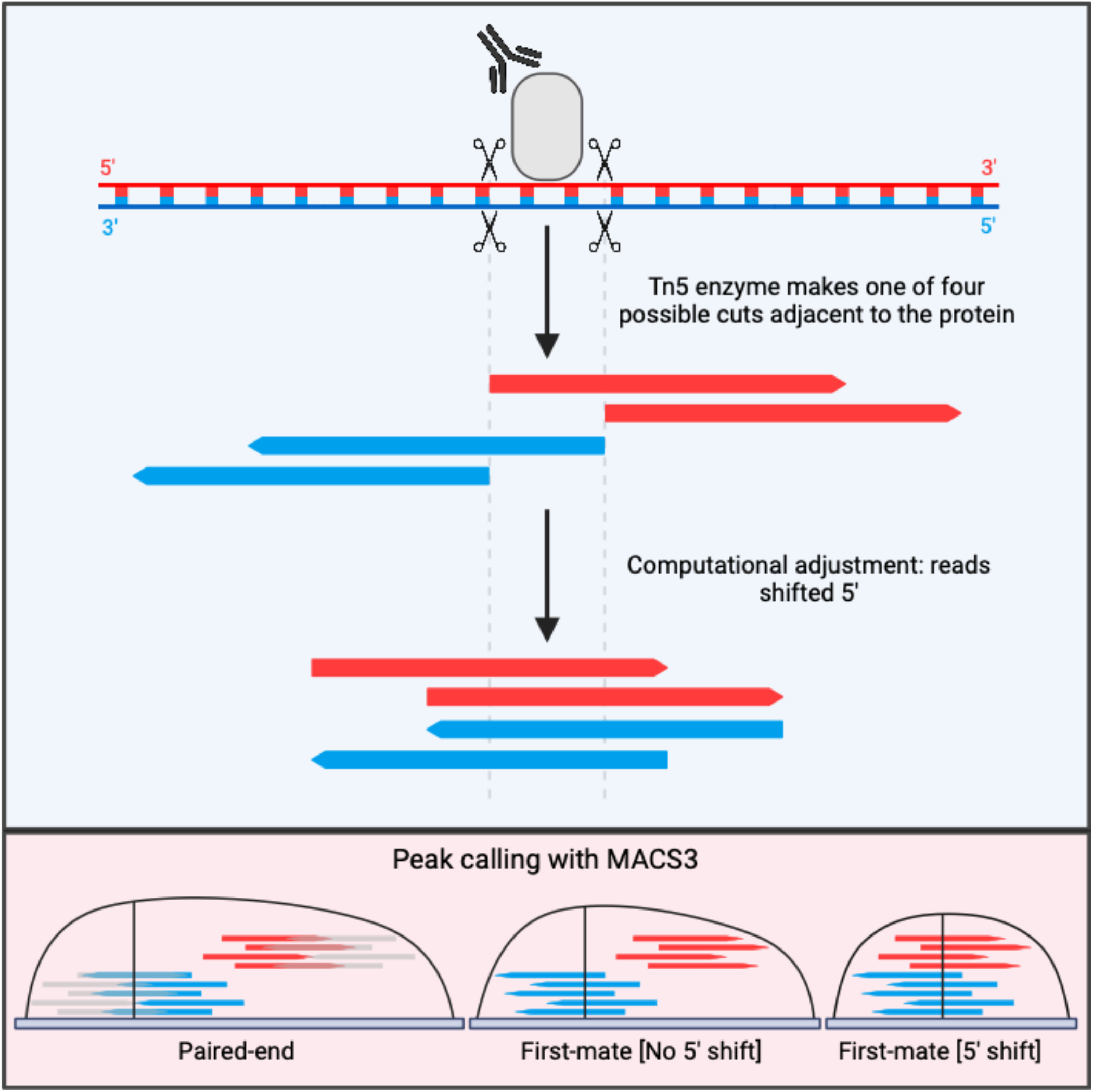
TIP-seq reads should be centred on their 5’ ends to reduce peak widths and the bias in summit positions. The pA-Tn5 enzyme makes one of four possible cuts adjacent to the protein of interest. Top: Paired-end reads are generated, but only the first read is shown here. The first base pair of each read represents the pA-Tn5 cut site, with sequencing occurring in the 5’ to 3’ direction. Due to the anti-parallel nature of DNA, sequence reads extend 3’ away from the cut site leading to wider peaks. Bottom: Paired-end sequencing further increases peak width (bottom left), while an observed preference for read generation on one of the strands biases the peak summit towards this strand (bottom left and centre). Using only the first reads (first mate) centred on their 5’ ends produces narrower peaks with unbiased summits (bottom right).

To assess the predictive power of peaks, we considered several possible read-outs. Past work, including that from our lab, has used ENCODE ChIP-seq peaks as benchmarks to assess the performance of Tn5-based methods (Hu et al., 2023; Meers et al., 2019; Yashar et al., 2022). However, there is no data to suggest that ChIP-seq outperforms other methods in detecting true protein binding. In addition, due to the unspecific fragmentation of DNA in ChIP-seq, reads belonging to one peak can span a relatively large area, while protein binding typically covers between 10-20 nucleotides. ChIP-seq alone therefore does not lend itself very well to assessing, e.g. the effect on peak width. However, when profiling transcription factors with known motifs, motif enrichment can serve as an objective metric to assess the quality of the profiling experiment (Kim et al., 2007). We propose that a TF peak set can be considered better if it (a) contains greater motif enrichment and (b) the peak summits are closer to those motifs.

### Single-read peak calling reduces peak widths

To test whether read centring and first read filtering improves Tip-seq data analysis, we used previously published paired-end CTCF Tip-seq data (∼3 Million reads), as well as in-house generated CREB1 Tip-seq data (∼6.5 Million reads) starting from raw read BAM files. All data is derived from paired-end sequencing, as this is usually more cost effective, and can improve mapping accuracy.

Peaks were called using MACS either using both or only the first read. As expected, using only the first read reduced the peak width to 55% and 58% for CTCF and CREB1, respectively (Table 1). To our surprise, despite reducing the data by half, the number of peaks from first reads only increased to 124% (CTCF) and 121% (CREB1) (Table 1), suggesting that the data reduction does not interfere with sensitivity. This is likely because paired-end peak calling simply increases the fragment length rather than doubling the number of fragments (Fig. 1).

**Table 1.** CTCF and CREB1 peak metrics under three peak calling settings. % refers to paired-end results.

|  | Paired-end | First read, no shift | First read, 5'-shift |
| --- | --- | --- | --- |
| **CTCF** |  |  |  |
| Peak number | 37,407 | 46,283 (124%) | 49,236 (132%) |
| Mean peak width (bp) | 536 | 293 (55%) | 231 (43%) |
| Mean distance to nearest motif (bp) | 99 | 62 (63%) | 31 (31%) |
| Median distance to nearest motif (bp) | 52 | 48 (92%) | 18 (35%) |
| Motif enrichment (primary / control) in %; e-values | 27.87 / 4.47;<br>4.50e-1796 | 25.18 / 2.88;<br>1.64e-2333 | 25.49 / 2.47;<br>1.15e-2686 |
| **CREB1** |  |  |  |
| Peak number | 46,371 | 56,452 (122%) | 60,179 (130%) |
| Mean peak width (bp) | 551 | 320 (58 %) | 286 (52%) |
| Mean distance to nearest motif (bp) | 167 | 98 (59%) | 82 (49%) |
| Median distance to nearest motif (bp) | 114 | 68 (60%) | 45 (39%) |
| Motif enrichment (primary / control) in %; e-values | 17.71 / 7.71;<br>9.02e-463 | 14.13 / 6.05;<br>5.21e-450 | 14.01 / 5.58;<br>2.20e-540 |

### Centring the 5’ ends of reads reduces summit-to-motif distance

In Tip-seq, and other protein-bound DNA profiling methods using linear amplification, the original cut site will be at the 5’ end of a read either upstream or downstream of the protein of interest. The 5’ end of the first read therefore represents the closest known point to the target, while the read itself extends downstream towards the 3’ end (Fig. 1), i.e. the 3’ bias. We hypothesise that centring peaks on cut sites by in-silico shifting upstream, as is suggested for DNase-seq (Zhang et al., 2008), would further reduce peak width. In addition, we noticed a bias of the peak summits towards the strand with more reads (Fig. 1), which is further aggravated by the 3’ bias. Reducing this bias through 5’ shifting should therefore improve peak accuracy. MACS summits detect the nucleotide with the highest read pile-up and are designed to detect the absolute binding sites. We can therefore use the offset between summits and binding motifs to measure peak accuracy.

While first-read peaks are narrower, we observed a substantial displacement between MACS peak summits and motif locations, resulting in a bimodal distribution peaking adjacent to but not on the motif itself, as also seen with paired-end data (Fig. 2). We suspect that this is due to the combination of an uneven distribution of reads across both DNA strands and the 3’ bias, as detailed in Fig. 1, skewing the summits towards the strand with greater read density and further diluting the signal. To test this, we shifted all reads 5’ by *½ ′ fragment size*, thus centring reads on the original cut sites (Fig. 1). This shift further reduced the average peak width for CTCF and CREB1 to 231 bp and 286 bp, respectively (Table 1). In addition, we obtained even more peaks compared to first-read peak calling without shift (Table 1), suggesting an increase in sensitivity. Importantly, the 5’ shift substantially lowered the average summit to motif distance to less than half compared to paired-end sequencing for both proteins (Table 1) and achieved a unimodal distribution of peak summits centred exactly on the motif (Fig. 2).

**Figure 2.**
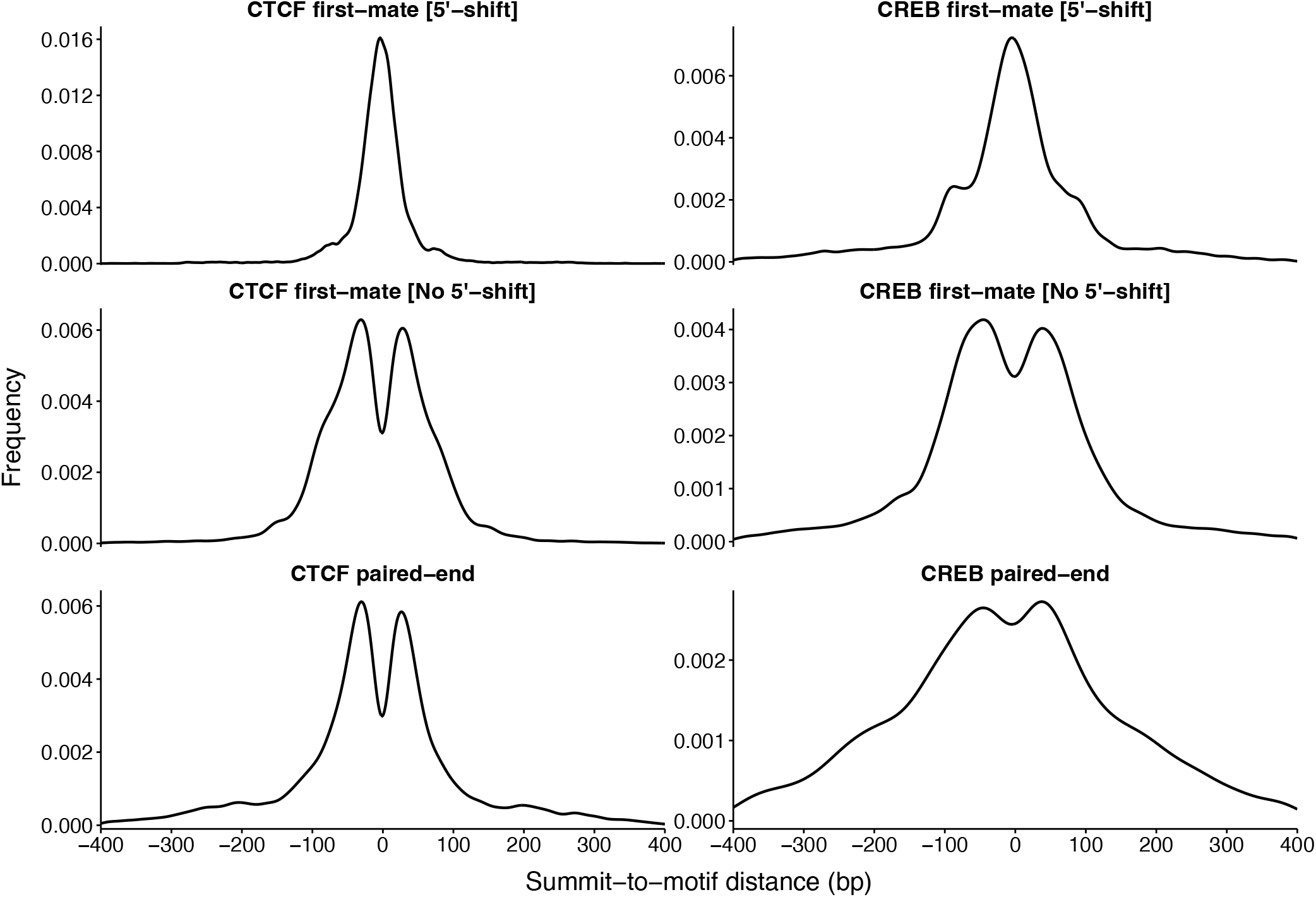
TIP-seq peaks are centred on transcription factor binding sites when MACS is run on read 1, 5’-shifted reads. Distribution of peak summits relative to CTCF and CREB motifs discovered using FIMO from the MEME suite. The plots compare three different peak calling methods: first-read [5’-shift] (top), first-read [no 5’-shift] (middle), and paired-end mode (bottom). The frequency of summit-to-motif distances (bp) indicates that read 1, 5’-shifted peaks (top row) show a clear centralisation around the motif for both CTCF (left column) and CREB (right column), compared to the other methods.

**Figure 3.**
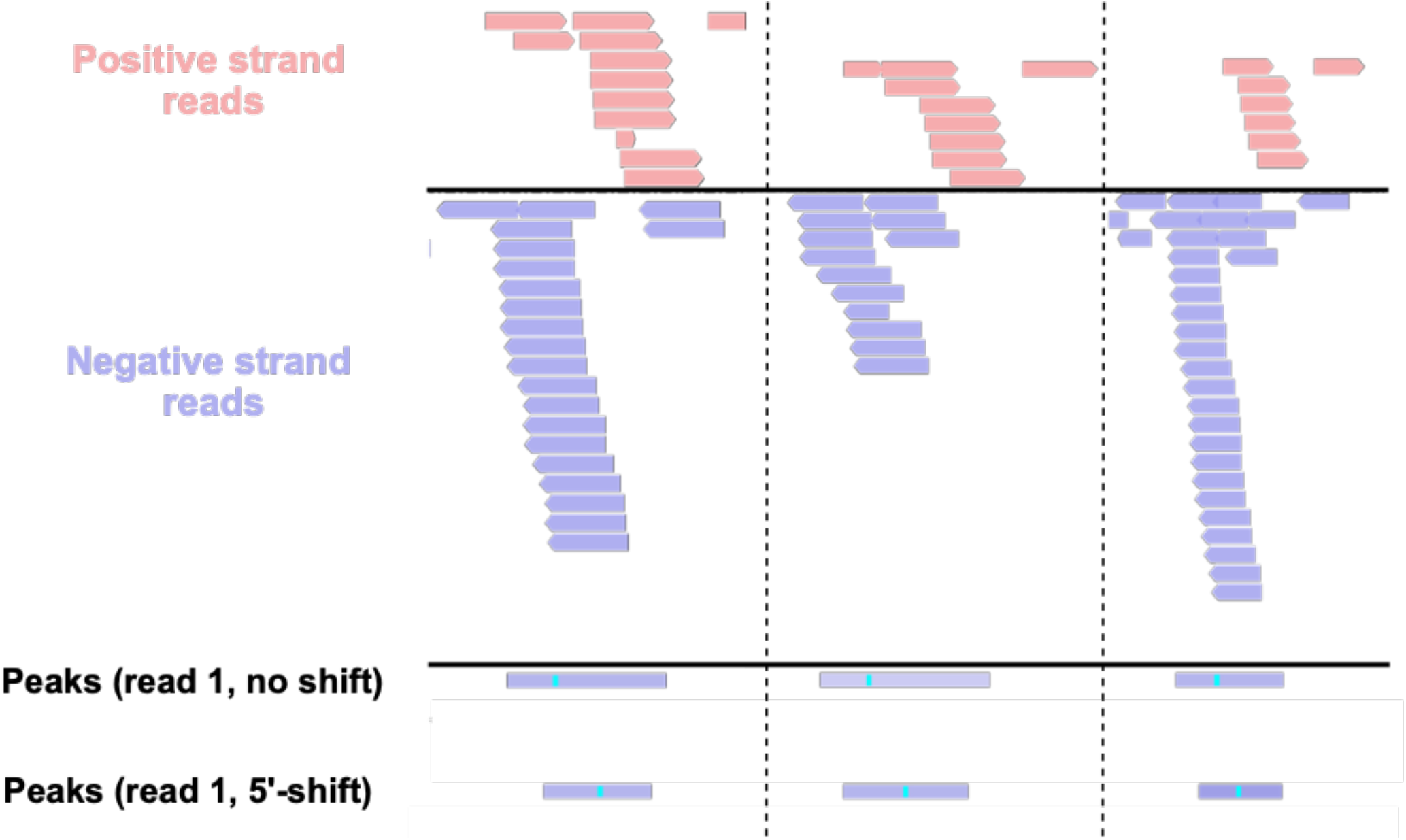
Real-world examples of TIP-seq data demonstrating the need to centre all reads on their 5’ ends. Each of the three columns shows an example of a peak from a real TIP-seq dataset, visualised using IGV. Only the first read of each read pair is shown. The red reads are aligned to the positive strand, and the blue reads are aligned to the negative strand. Whether a read is on the positive/negative strand is random: in theory, there should not be a bias, but in real-world data, there generally is. The misalignment in positive and negative strand reads can be seen in each column. If the reads are not shifted prior to peak calling, this skews the summit positions towards the strand with the most reads. The peaks that are called from the data, with and without read shifting, are shown along the bottom. The vertical cyan line within each peak denotes the summit. We expect the target protein to bind at the midpoint between the positive and negative strand reads, which should be reflected in the summit placement. In all examples, the read1, no 5’-shift peak summits are biased toward the strand with the most reads, whereas the read 1, 5’-shift summits are correctly centred on the midpoint.

### First-read, 5’-shifted peaks are enriched for motifs

To test the specificity of the first-read, 5’-shifted peaks, we calculated the relative enrichment of the motif in these peaks compared to the unshifted method using AME from the MEME suite. 11,652 first read, unshifted CTCF peaks contained the motif (25.18%) relative to 1,332 control sequences (2.88%) (e-value = 1.64e-2333). Similarly, 12,551 first-read, 5’-shifted CTCF peaks contained the motif (25.49%) compared to 1,217 control sequences (2.47%) (e-value = 1.15e-2686). Of the 3,640 unique peaks not captured by the first-read, non-shift method, 10.27% were enriched for the motif relative to 1.43% of control sequences (e-value = 4.60e-62), highlighting that the additional peaks captured are genuine. Results were similar for CREB1 (Table 1). Note that while the motif enrichment generally decreases as a function of peak width, the significance scores corrected for peak width (e-values) improve substantially for both targets (Table 1), proving superiority for motif identification with our optimised analysis parameters.

### *De novo* motif discovery is more effective on narrowed peaks

*De novo* motif discovery typically follows peak calling and aims to identify short nucleotide sequences (i.e. motifs) underlying TF binding. Motifs are identified based on their enrichment within the peak set. Narrow peaks reduce the amount of surrounding, non-relevant DNA sequences and should thereby increase the signal-to-noise ratio, while lowering computational runtime. Given that the summits of the first-read, 5’-shifted peaks are closely aligned with motifs, we hypothesised that entire peaks are not necessary for motif detection. To test this, we varied CTCF peak widths from 50-300bp in intervals of 50bp, centred on peak summits and performed *de novo* motif discovery using STREME.

Our results indicate that *de novo* motif discovery is most effective on summit-centred, read 1, 5’-shifted peaks, which discovered the CTCF motif as the top hit at all peak widths. For this modality the enrichment score quickly plateaued starting at 100bp (48.6%, runtime = 8 minutes; Fig. 4), with the highest enrichment at 200bp (54.5%) with a runtime of 16 minutes and lowest at 50bp (29.1%, runtime = 4 minutes). In contrast, neither the paired-end peaks nor the read 1, no-shift peaks detected the CTCF motif at a peak width of 50bp. The paired-end peaks reached their highest enrichment at 300bp (52.3%, runtime = 21 minutes), while the read 1, no-shift peaks peaked at 250bp (54.2%, runtime = 18 minutes). Taken together, the optimised first-read, 5’ shifted peak calling parameters improve motif discovery through peak narrowing and motif centring, while reducing the computational resources needed.

**Figure 4.**
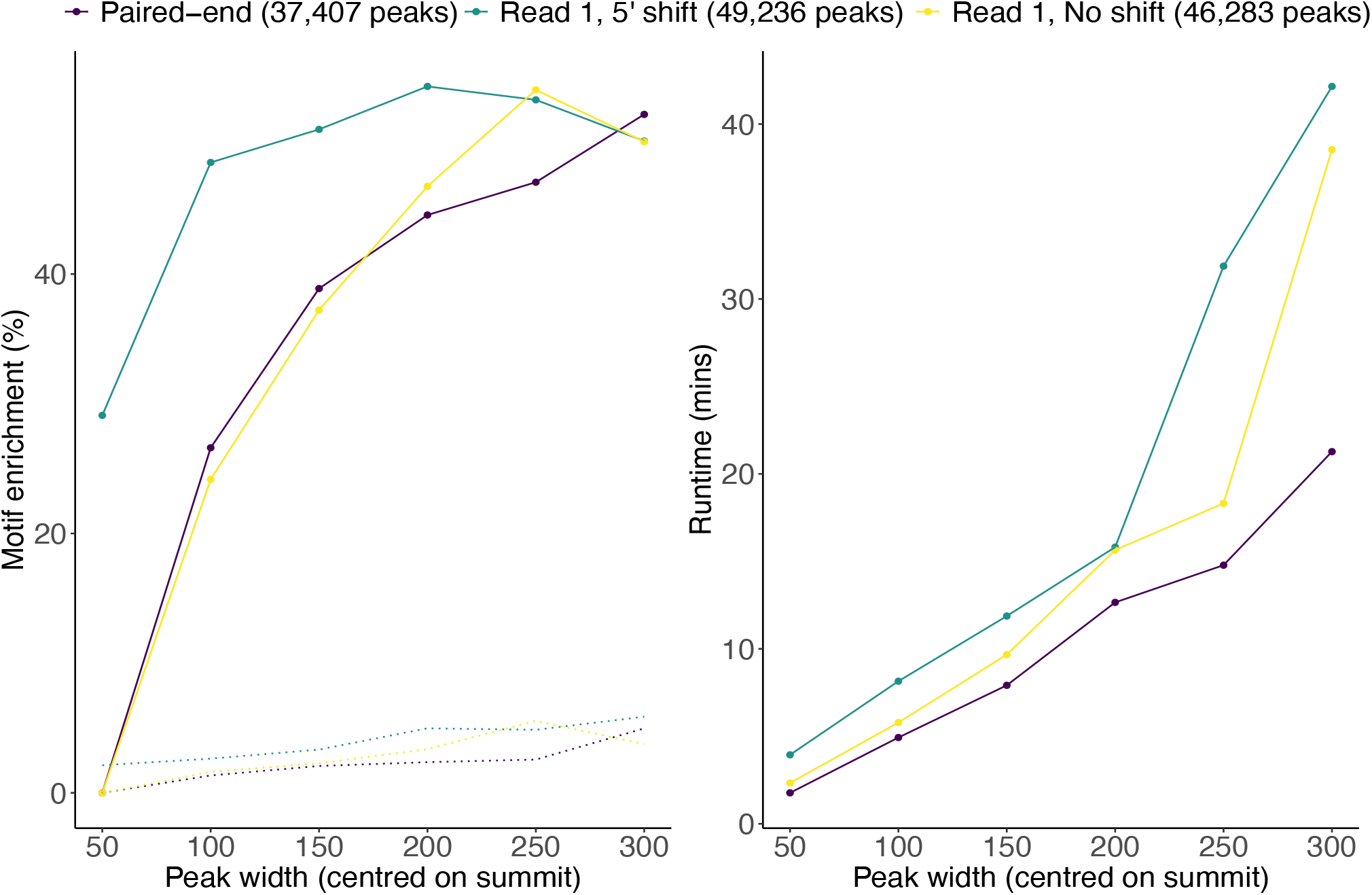
Higher motif enrichment at lower peak widths for read 1, 5’-shifted peaks. We evaluated the e+ect that di+erent peak calling settings had on de novo motif discovery using % motif enrichment. Three di+erent peak sets were generated using MACS3: (1) a peak set using the paired-end mode, (2) a peak set using only read 1 of each read pair, and (3) a peak set using only read 1 with the reads shifted to centre them on their 5’ ends. De novo motif discovery was then performed on filtered versions of each peak set. Peaks were centred on their summits and extended to widths of 50, 100, 150, 200, 250, and 300 bp. **(A)** Peaks called in the read 1, 5’-shift setting detected the known motif in a higher percentage of peaks; this was particularly pronounced when only narrow peaks around the summits were used **(B)** De novo motif discovery runtime scales with peak width. When balancing % motif enrichment and runtime, the read 1, 5’-shifted peaks facilitate quicker de novo motif discovery.

## Discussion

Understanding the technical nuances of DNA enrichment assays is crucial for effective peak calling. Given the status of MACS peak caller as industry standard, users require a thorough understanding to determine the optimal settings, particularly when a method is novel. Our study leverages the precision of TIP-seq’s single-adaptor insertion through pA-Tn5 transposase to optimise MACS peak calling. By analysing the first read of read pairs only and incorporating a 5’ shift to counteract bias from frequent strand preference for adaptor insertion and amplification in the 3’ direction, we have narrowed average peak widths and reduced the distance between peak summits and their respective motifs. Due to the similarities between TIP-seq and CUT&Tag, we expect this logic to also apply to some CUT&Tag-based protocols, especially Nano-CT, which also uses linear amplification. NTT-seq and uCoTarget, where the first read is defined through a primary antibody and the second read through a secondary antibody, would likely benefit from the 5’ shift to centre on primary cut sites, while we expect a smaller effect on peak width compared to Tip-seq, given that both antibodies should be spatially close to the target.

An interesting aspect of our peak calling strategy is the increased number of peaks. Given the high performance in motif enrichment and discovery, we expect that the additional peaks are genuine, suggesting that our analysis pipeline is more sensitive compared to paired-end peak calling. Although paired-end sequencing is generally more economical when outsourcing and can improve the alignment of reads to the genome, we argue that single-end sequencing would be sufficient to obtain high-quality data for Tip-seq.

For CUT&Tag based workflows where adaptors for both reads are inserted simultaneously through target-bound pA-Tn5, 5’ shifting of paired-end sequencing is not currently possible with MACS. We suggest instead to treat both reads as first reads after removal of PCR duplicates, since both are derived from the same targeted insertion event. This should double the number of reads for peak calling and allow 5’ shifting. Additional work using peak calling on transcription factors with motif enrichment should be used to test this analysis strategy for CUT&Tag data.

Comparing different peak calling approaches requires a robust benchmarking metric. Commonly, ENCODE ChIP-seq peaks are used for this purpose, with the assumption that they denote true positives. Ideally, we would use an unbiased benchmark independent of peaks and this has motivated the use of motifs. Integrating motif information with peaks has allowed us to identify optimal peak calling parameters. Specifically, we have used the average distance between peak summits and their nearest motifs as an indicator of peak precision. We can do this because peak summits are proxies for TF binding sites. Using this, we have been able to confirm that TIP-seq peak calling is most effective when reads are centred on their 5’ ends prior to peak calling. This metric would prove helpful when developing new peak callers for specific assays.

While using motifs to benchmark has potential, it is important to note that the values for motif enrichment calculated by programs like AME and STREME are no absolute measure and are highly sensitive to the specific motif and the significance threshold. However, by observing relative changes in motif enrichment across different parameter settings, we have been able to draw meaningful conclusions about the effectiveness of different peak calling parameters. Therefore, while the absolute enrichment values should be interpreted with caution, they remain valuable for comparative analyses.

Another consideration is the use of multiple replicates for consensus peak calling to create a high-confidence set of peaks. Irreproducible Discovery Rate (IDR) thresholding is a popular method for this and is included in ENCODE’s ChIP-seq processing pipeline (Hitz et al., 2023; Q. Li et al., 2011). While here we only had access to single replicate TIP-seq data, it is likely that our motif enrichment values are higher on IDR-thresholded peaks due to the removal of non-reproducible, less specific peaks.

In conclusion, our optimised TIP-seq peak calling strategy demonstrates significant improvements in peak resolution, sensitivity, and accuracy as demonstrated by aligning peak summits with known motif locations. While acknowledging the limitations of single-replicate data and the variability in motif enrichment values, our approach offers a significant advancement in identifying true TF factor binding sites.

## Code availability

All code used in our analysis has been incorporated into an R package, *MotifPeeker*, which is available through GitHub (https://github.com/neurogenomics/MotifPeeker.git). This package is under CI/CD. A script which performs the specific analyses for this study is available on GitHub (https://github.com/neurogenomics/read1_peak_calling).

## Availability of data and materials

The CREB1 read data is available on the European Nucleotide Archive under study accession PRJEB79242. The CTCF read data was made available by Bartlett et al. (2021) and can be found in NCBI GEO under accession number GSE188512.

